# Macro-to-nano scale investigation of wall-plate joints in the acorn barnacle *Semibalanus balanoides*: correlative imaging, biological form and function, and bioinspiration

**DOI:** 10.1101/590158

**Authors:** R.L Mitchell, M. Coleman, P. Davies, L. North, E.C. Pope, C. Pleydell-Pearce, W. Harris, R. Johnston

## Abstract

Correlative imaging combines information from multiple modalities (physical-chemical-mechanical properties) at various length-scales (cm to nm) to understand complex biological materials across dimensions (2D-3D). Here, we have used numerous coupled systems: X-ray microscopy (XRM), scanning electron microscopy (SEM), electron backscatter diffraction (EBSD), optical light microscopy (LM), and focused-ion beam (FIB-SEM) microscopy to ascertain the microstructural and crystallographic properties of the wall-plate joints in the barnacle *Semibalanus balanoides*. The exoskeleton is composed of six interlocking wall-plates, and the interlocks between neighbouring plates (alae) allow barnacles to expand and grow whilst remaining sealed and structurally strong. Our results indicate that the ala contain functionally-graded orientations and microstructures in their crystallography, which has implications for naturally functioning microstructures, potential natural strengthening, and preferred oriented biomineralisation. Elongated grains at the outer edge of the ala are oriented perpendicularly to the contact surface, and the c-axis rotates with the radius of the ala. Additionally, we identify for the first time three-dimensional nano-scale ala pore networks revealing that the pores are only visible at the tip of the ala, and that pore thickening occurs on the inside (soft-bodied) edge of the plates. The pore networks appear to have the same orientation as the oriented crystallography, and we deduce that the pore networks are probably organic channels and pockets which are involved with the biomineralisation process. Understanding these multi-scale features contributes towards an understanding of the structural architecture in barnacles, but also their consideration for bioinspiration of human-made materials. The work demonstrates that correlative methods spanning different length-scales, dimensions and modes enable the extension of structure-property relationships in materials to form and function of organisms.

## 2. Introduction

Biomineralised organisms show an incredible diversity of complex microstructural forms and structure-property relationships [1–6]. A more complete realisation of these naturally-occurring structures provides not only a better understanding of an animal’s ecology[7–10], but also supports bioinspired development of human-made materials [11–16]. Acorn barnacles (order Sessilia) are sessile marine arthropods that often inhabit the high-energy intertidal zone and have adapted structurally, compositionally, and architecturally to challenging abiotic conditions, as well as the threat of diverse predators [17]. The calcareous exoskeleton (shell) of barnacles is well-studied structurally; for example the specific calcite crystal orientations in the operculum of *Balanus amphritrite* (= *Amphibalanus amphitrite*; [18,19]); the high mechanical strength and adhesive properties of the baseplate in *A. amphitrite, A. reticulatus* and *B. tintinnabulum* ([9,20–23]); the involvement of extracellular matrix molecules in exoskeleton biomineralisation in the giant barnacle *Austromegabalanus psittacus* [24]; and the structurally-sound nanomechanical properties of the exoskeleton of *A. reticulatus* [25]. However, little is understood about how macro-micro-nano-scale structures, particularly in the shell, are linked. Correlative imaging provides an opportunity to discover the multi-scale interactions and mechanisms involved in the structure of complex systems at varying length scales [26–29], and specifically for barnacles, provides an opportunity to correlate optical, analytical, structural, and mechanical information [30] for the first time. Here, we have coupled numerous systems at various length scales: X-ray microscopy, scanning electron microscopy, light microscopy, and focussed ion-beam microscopy to ascertain the macro-to-nanoscale structure, crystallographic orientation and mechanical properties of wall plate joints in the parietal exoskeleton of the barnacle *Semibalanus balanoides*.

*S. balanoides* is the commonest intertidal barnacle on British coastlines [31] commonly outcompeting other barnacle genera [7], although the structural properties of its shell are relatively poorly understood compared to other species (e.g., *B. amphritrite*), as are the morphological properties of the wall-plate joints, with just two previous studies outlining basic details [7,17]. The shell of *S. balanoides* comprises six interlocking joints, where the shell originates from within an existing organic cuticle. These joints are located in a particularly active and dynamic region of the barnacle shell and provide a waterproof seal and structural integrity in the face of extreme conditions of the physically harsh intertidal zone [8]. This work identifies specific macro-micro-nano features of the wall-plate joints in both 2D and 3D through connected correlative imaging and establishes how these features are linked at varying length scales. A greater understanding of how these complex structures function provides valuable biomechanical information for biologists as well as the broader bioinspiration topic.

## 3. Methods

### 3.1. Barnacle (Semibalanus balanoides) structure

Acorn barnacles are sessile organisms that attach to hard substrates via either a calcified base plate or an organic membrane [20], and biomineralisation of the calcareous shell is mediated by the mantle epithelium via secretion of a calcium matrix [32]. The conical-shaped exoskeleton is composed of four, six, or eight wall-plates depending on the species [25] which overlap at sutures, or joints (figure 1*a*); parts of the plate overlapping internally are called alae (‘wings’), and parts that overlap externally are called radii (‘rims’ [33]; figure 1*b*). The wall plates grow both upwards towards the apex and outwards as the internal soft-bodied organism grows inside [20]. As with other crustaceans, barnacles moult the chitinous exoskeleton surrounding their main body periodically to grow, but the calcareous shell is not shed during this process [33].

**Figure 1:**
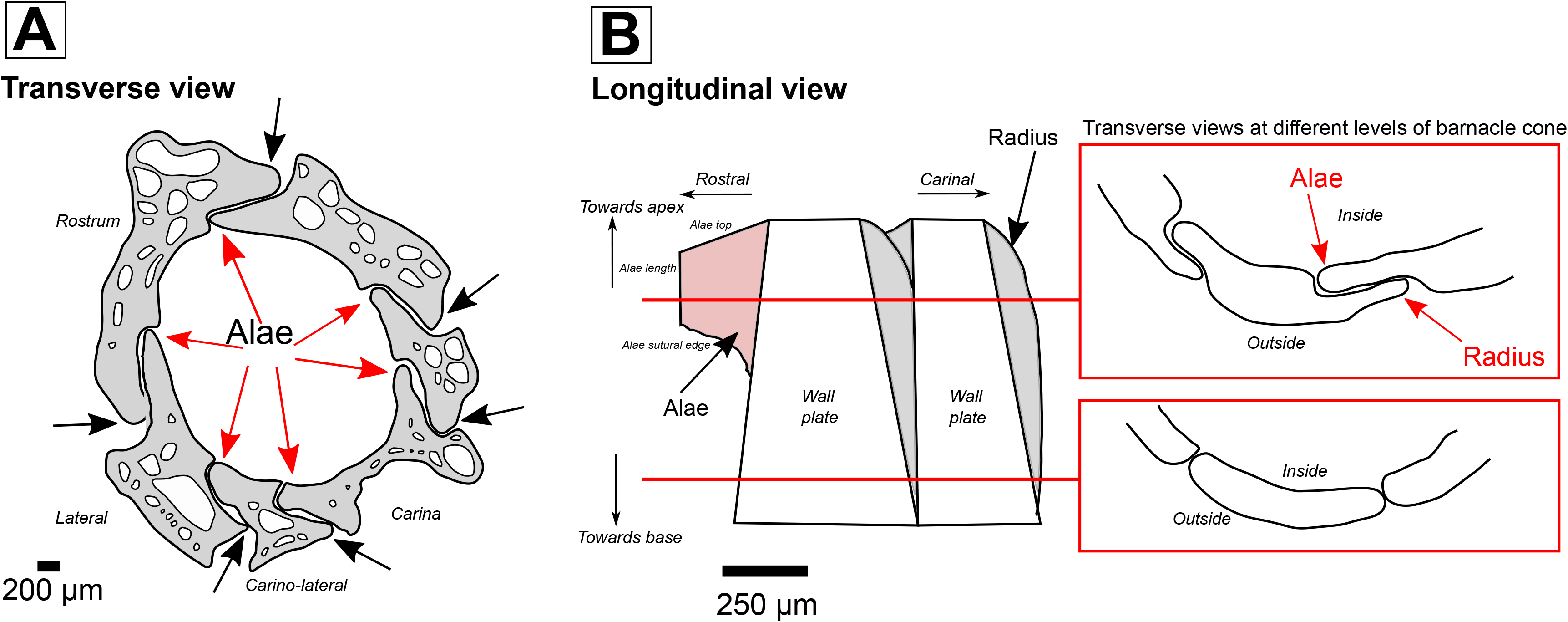
Morphological structure of the barnacle *Semibalanus balanoides. (a)* Transverse view of the six wall plates composing the barnacle conical structure. Alae between adjacent wall plates are highlighted by red arrows, radius on neighbouring plates by black. *(b)* Longitudinal internal view through adjacent wall plates. Inserts illustrate morphological differences of the ala at different points of the interlock. B adapted from [13].

### 3.2. Sample and preparation

Barnacle specimens were collected from the intertidal region at Bracelet Bay, Swansea, UK (51.5660° N, 3.9801° W). Samples were subsequently vacuum impregnated into a 32mm-wide resin block and ground and polished to reveal a transverse section. The sample surface was coated with a 10nm layer of carbon to ensure sample conductivity in the SEM and FIB-SEM. Individual plates were also detached from the exoskeleton and attached with adhesive to wooden pins for imaging using XRM. All preparation, subsequent analysis and imaging occurred within the Advanced Imaging of Materials (AIM) Facility within the College of Engineering at Swansea University (UK).

### 3.3. Imaging and analysis

#### 3.3.1. Light microscopy (LM) and Scanning Electron Microscopy (SEM)

LM and SEM were used to obtain general 2D information on barnacle morphology. LM images were obtained using a Zeiss SmartZoom and a Zeiss Observer Z1M inverted metallographic microscope. SEM images were collected on a Carl Zeiss EVO LS25 with a backscatter detector at 15kV, 750pA, and a working distance of 10mm. As well as the carbon coating, copper tape and silver paint were added to the sample surface to aid charge dissipation.

#### 3.3.2. Electron Backscatter Diffraction (EBSD)

A JEOL 7800F FEGSEM and a NordlysNano EBSD detector controlled via Aztec (Oxford Instruments) software were used to obtain crystallographic information. The phase selected for EBSD indexing was Calcite [34] and patterns were collected at 15kV with a step size of 0.2µm. A relatively high number of frames (5 frames per pattern) were collected using 4 × 4 binning to give a camera pixel resolution of 336 × 256 pixels and a speed of 8 Hz.

#### 3.3.3. X-ray micro Computed Tomography/Microscopy (XRM)

A Zeiss Xradia Versa 520 (Carl Zeiss XRM, Pleasanton, CA, USA) was used to carry out high resolution X-ray microscopy (XRM) non-destructive imaging; this was achieved using a CCD detector system with scintillator-coupled visible light optics and a tungsten transmission target. Initial scans of the barnacle region block were undertaken with an X-ray tube voltage of 130 kV and a tube current of 89 μA, and an exposure of 4000 ms. A total of 1601 projections were collected. A filter (LE4) was used to filter out lower energy X-rays, and an objective lens giving an optical magnification of 4 was selected with binning set to 2, producing an isotropic voxel (3D pixel) size of 3.45μm. The tomograms were reconstructed from 2D projections using a Zeiss Microscopy commercial software package (XMReconstructor), and an automatically generated cone-beam reconstruction algorithm based on filtered back-projection. Individual plates were also scanned (not in the resin block); these were collected using the 4X objective lens at 60kV and 84µA, with an exposure time of 12000 ms and a resulting (isotropic voxel size) of 0.5 µm. A filter (LE1) was used to filter out low energy X-rays, and 1601 projections were collected. The scout and zoom methodology was used to create high resolution regions of interest within the sutures.

### 3.4. Correlative Microscopy (Zeiss Microscopy Atlas 5/3D)

Targeted navigation to regions of interest was achieved using Zeiss Microscopy correlative Atlas 5 (3D) software package on the Zeiss Crossbeam 540 FIB-SEM. This method enables a live 2D SEM view to be combined with other data and information from previous sessions or relevant characterisation techniques on the same area or volume of interest; this is achieved by importing and aligning other 2D datasets (e.g., LM images, EBSD maps) and 3D data (XRM stacks) to accurately correlate and locate regions of interest for further nano-scale imaging and characterisation (figure 2). Initial overlay is achieved by manually aligning the live SEM image with the imported data, and ‘locking in’ the imported data to the current SEM coordinate system. This correlative microscopy approach is especially useful when regions of interest may be internally located within a subsurface area of the specimen, and allows samples to be accurately studied at varying length-scales by combining information from multi-modal sources.

**Figure 2:**
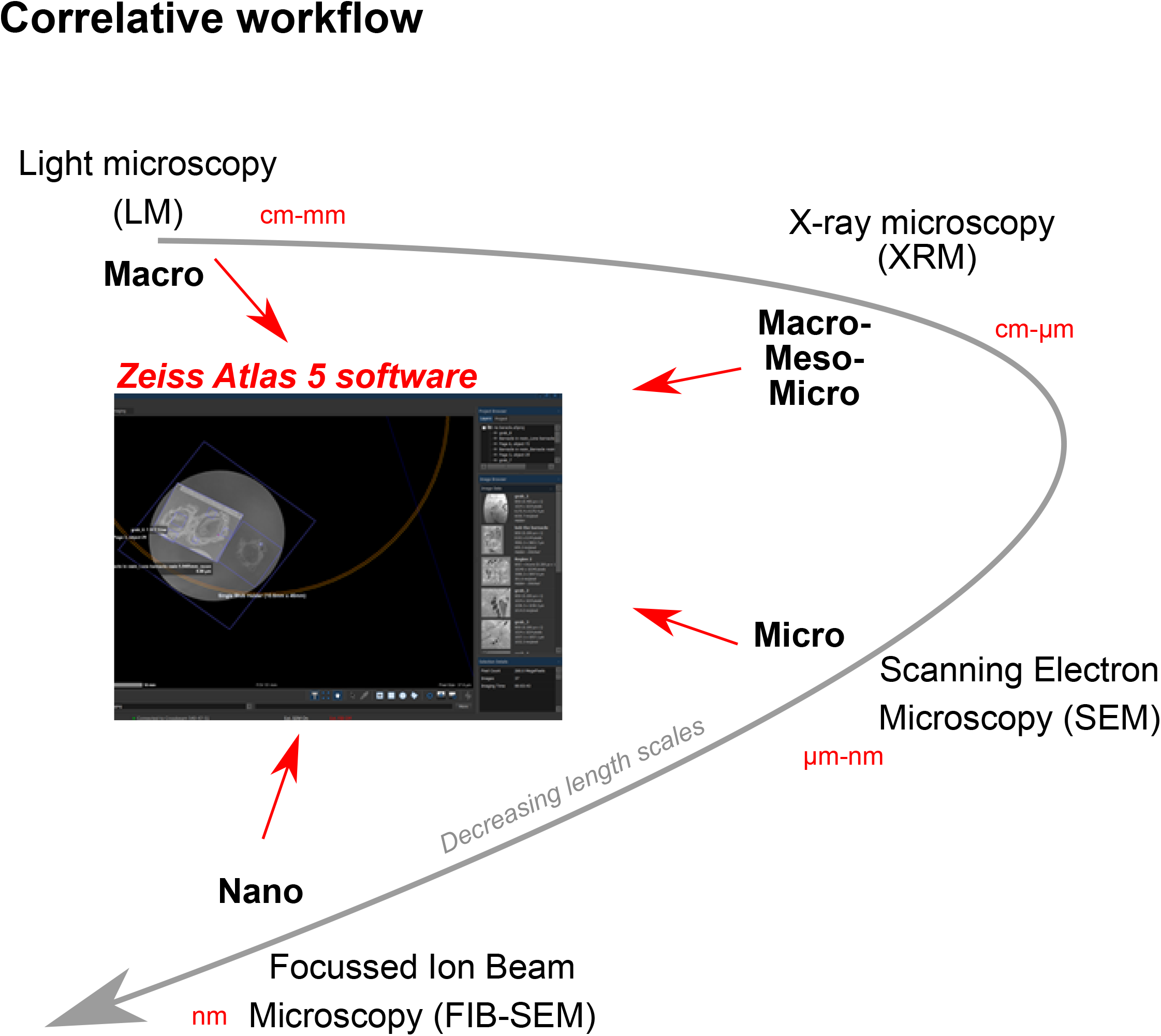
Schematic of the multi-modal and multi-scale correlative workflow utilising LM, XRM, SEM, AND FIB-SEM. These techniques can be correlated using Zeiss Microscopy Atlas 5 (3D) software.

#### 3.4.1. Focussed Ion Beam Microscopy (FIB-SEM)

Specific regions of interest in the barnacle shell were studied using a Zeiss Crossbeam 540 Focussed Ion Beam Scanning Electron Microscope (FIB-SEM, Gallium source; Carl Zeiss Microscopy, Oberkocken, Germany). The sample stage was tilted to 54° to allow the sample to be perpendicular to the FIB column; the ion beam energy was 30kV in all cases. The FIB and SEM beams are then aligned at 5mm working distance at the co-incidence point. Within the Atlas 5 (3D) correlative workspace it is possible to identify regions of interest for further study, and then with the same interface prepare and collect 3D nanotomographic volumes (figure 3). A template is set up over the region of interest which outlines the numerous steps in the milling process (figure 3 *a*). A 10 × 10µm platinum layer was deposited using a gas injection system and the 700pA FIB probe; this is to protect the ROI sample surface from damage during the milling process. 3D tracking marks (which enable automatic alignment and drift correction during an automated run) are milled onto the first platinum pad using the 50 pA FIB probe, and then a second platinum pad is deposited on top (again at 700pA) creating a ‘sandwich’ of protection and alignment layers (figure 3 *b*). A trench is then milled using the 7nA probe to create a cross sectional surface through the region of interest to a depth of ~15 µm (figures 3 *b, c*). The cross-sectional surface of the trench is polished using the 700pA probe. Once the sample preparation is complete, automated tomographic milling and slice generation can take place (figure 3 *c*). The run is set so the length of the protected platinum pad is milled to create a 3D volume. Each slice (10nm thickness) is milled by the 1.5nA probe using the FIB and simultaneously imaged by the SEM; parameters for image acquisition with the SESI detector include 1.8kV, 300pA, 10 µs dwell time and a 12nm pixel size. Once the run has completed overnight (~8 hours), the slice images are reconstructed to create a 3D volume (figure 3 *d*), and segmented and visualised via other specialised tomographic software (e.g. FEI Avizo, Hillsboro, USA; ORS Dragonfly, Montreal, Canada). Quantified data can be found in *supplementary material 1*.

**Figure 3:**
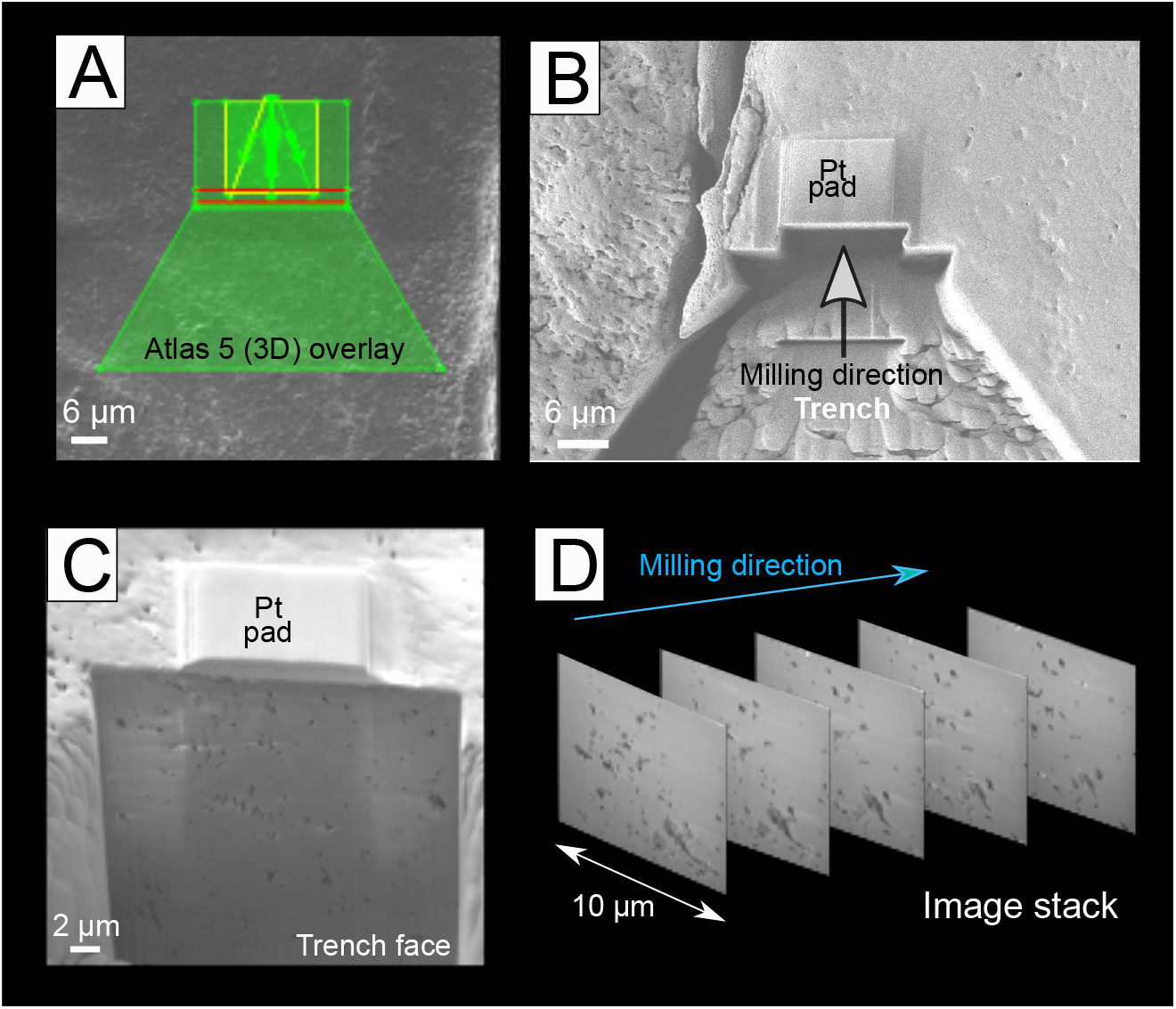
Stages of the FIB-SEM automated milling process using Zeiss Microscopy Atlas 5 (3D). *(a)* An overlay is created for each part of the milling preparation*. (b)* Once a platinum pad has been deposited over an initial platinum pad and the milled reference marks creating a ‘sandwich’, a trench is milled to reveal a cross section face *(c). (d)* Acquired images can then be stacked together to create a tomographic volume.

## 4. Results

### 4.1. 2D ala morphology and crystallography (SEM and EBSD)

SEM reveals the micro-scale 2D morphology of the barnacle (and specifically the alae; figure 4). Alae generally have rounded tips and slot into a groove in the neighbouring plate with organic material separating the two plates (figure 4 *a-d*). The microstructure in the 40-70 µm closest to the tip of the ala appears to have a different morphological texture and more voids than other alae regions and the opposing plate (figures 4 *c, d*). The voids are of two types; transverse banding which is parallel to the ala tip orientation, and elongated grooves/channels, which are perpendicular to this (figure 4 *c, d*). The elongated grooves/channels and transverse banding appear to be of varying size, shape, elongation and thickness (figures 4 *c, d*) and may represent pore networks. In contrast, the neighbouring plate and the area behind the ala tip appear smooth and featureless (figure 4 *c, d*).

**Figure 4:**
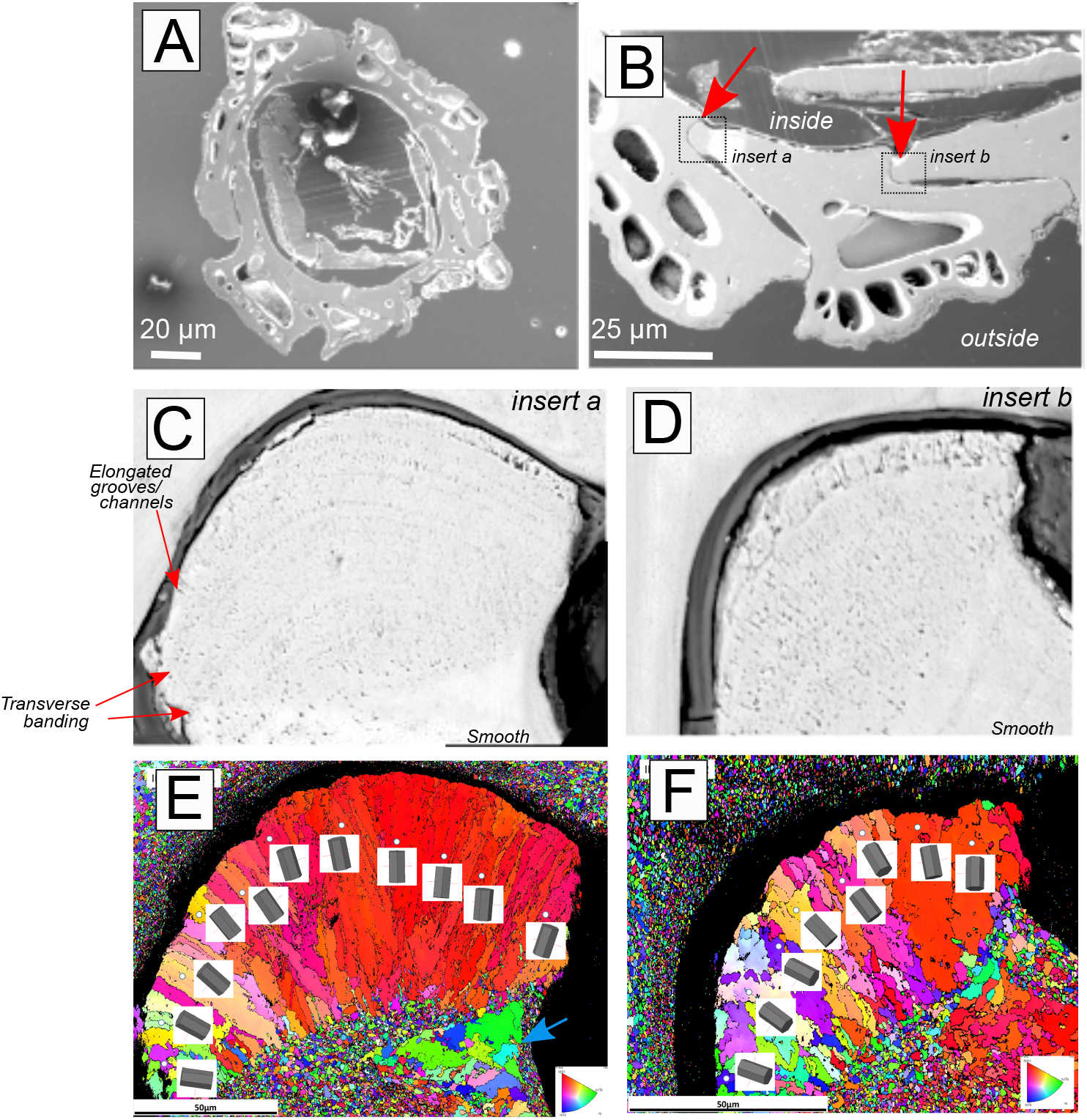
SEM imaging and EBSD crystallography of the barnacle and ala. *(a)* Scanning electron microscope (SEM) image of an individual barnacle in transverse section. *(b)* View of three interlocking plates and ala (red arrows). *(c-d)* Close up of two ala (insert boxes in *b*), revealing micro-structure transverse banding and perpendicular elongated grooves/channels at the tip. *(e-f)* Electron backscattered diffraction (EBSD) maps of ala in *c-d*, illustrating elongated grain orientations at the tip of the ala, and granular grains behind the tip and on the adjacent plate. Blue arrow illustrates inside edge coarse grains. Elongated grains appear to correlate with porous area of ala. Scales in e and f correspond to c and d, respectively.

EBSD inverse pole figure maps of the ala tips show a microstructure with a highly-segregated bimodal grain size (figures 4 *e, f*). The grains at the tip of the ala closest to the plate joint are elongated and radiate 50 − 70µm downward into the ala structure perpendicular to the curve of the join (figures 4 *e, f*). The individual 3D hexagonal crystal diagrams for each elongated grain show that in each case the c-axis [0001] is parallel to the long axis of the grain and perpendicular to the line of the join of the plates (figures 4 *e, f*). The grains in the adjacent area below the elongated grains, and in the adjoining upper plate, are around 10-20 times smaller at 3 - 5µm, and have a more equiaxed structure with no obvious texture. In both EBSD images there are also regions of coarser grains within the equiaxed areas behind the ala tip on the inside-facing edge of the ala (blue arrow; figure 4 *e*), however these are not elongate or organised like those in the tip (figure 4 *e, f*).

### 4.2. 3D ala morphology and porous networks (XRM and analysis)

We have reconstructed the entire barnacle in 3D (figure 5 *a*; *supplementary material 2*) as well as individual plates (figure 5 *b*) illustrating morphological variations in the ala through the length of the exoskeleton. We observe the protruding ‘tab’ of the ala towards the apex where it slots in to the neighbouring plate (figure 5 *a ii*); in 2D image slices (figure 5 *a iii, iv*), the ala appears as a finger-like protrusion with a rounded tip. Towards the base and the ala sutural edge, the ala recesses, and creates a flat junction with the neighbouring plate (figure 5 *a ii*); in 2D the ala appears more angular and has an almost square tip (figure 5 *a ii, iv*). In addition to the 3D morphological change in the ala through its length, we also observe networks of elongate channels, grooves and bands that are also visible in the 2D surface imaging (figure 4 *c, d*). We propose that these are related to the pore networks observed in figure 5. Pores appear black in 2D stack images because they exhibit a lower X-ray attenuation compared with the surrounding calcium carbonate exoskeleton (figure 5 *a iii, iv*). The pores appear to ‘fan’ perpendicularly to the ala edge, the same as the textures in 2D (figures 4 *c, d*). The micropores are only found at the tip and are not present in other areas of the ala. Pores also change shape, orientation and location through the length of the ala; towards the apex they fan around the entire ala tip (figure 5 *a iii*), however towards the ala sutural edge they are on one, inner side only (figure 5 *a iv*).

**Figure 5:**
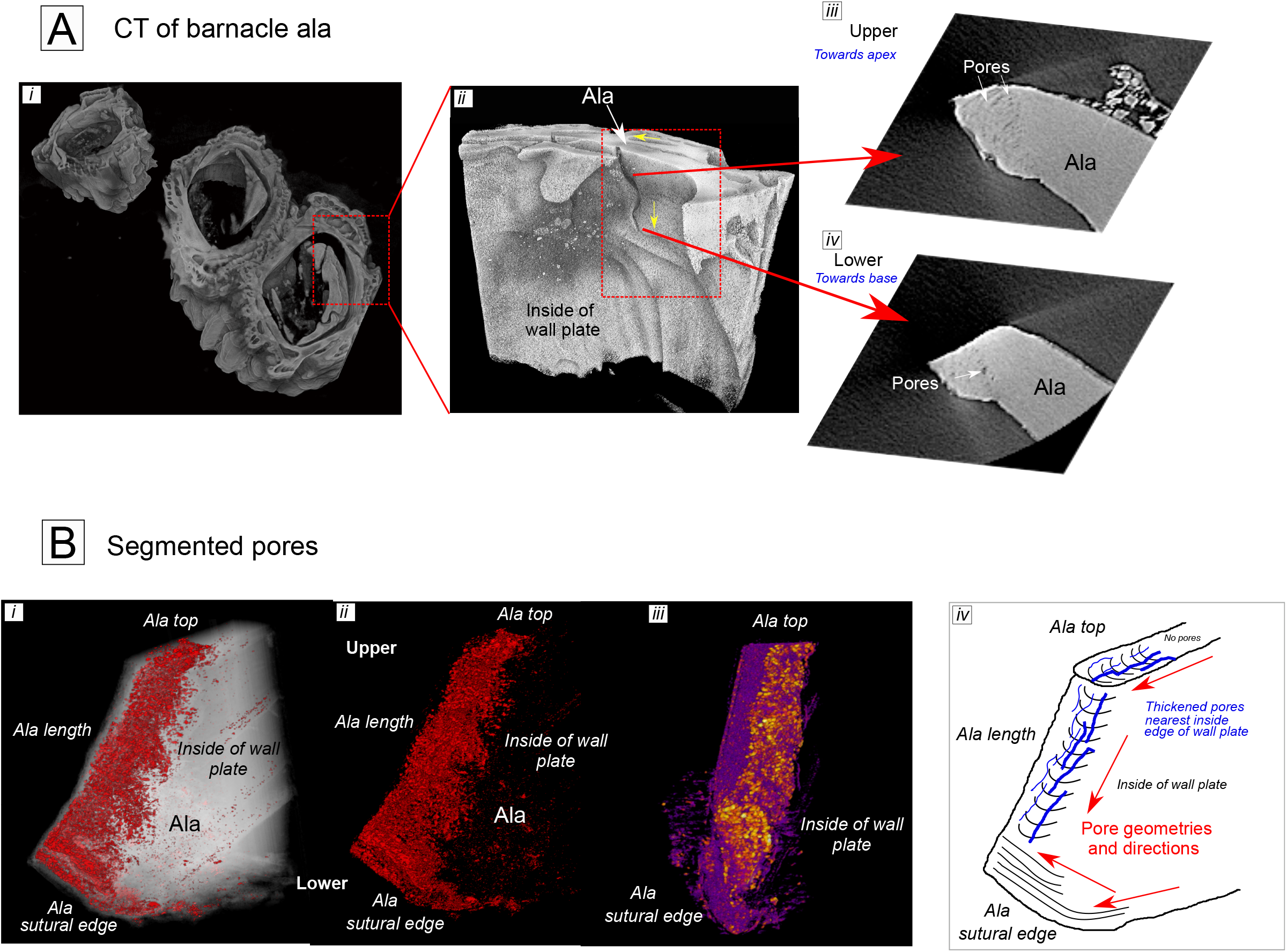
XRM of the barnacle. *(a, i)* 3D XRM image of the barnacle. *(ii)* XRM reveals changes in the morphology of the ala through its length. Also indicated are the directions upon which the ala meets the neighbouring plate (yellow arrows). Also identified are pore networks, and how these change through the ala length (*iii, iv*). *(b)* Segmented XRM ala pores (*i-iii*). From local thickness analysis in Fiji, pores appear to be thickest on the inside-edge of the plate, nearest the soft body of the organism. Purple = thin, yellow = thick. *(iv)* Simplified illustration showing the change in pore geometry through the length of the ala; the pores (blue lines) are parallel to the direction of the ala which continues down the ala length. Once at the ala sutural edge, the pores change direction, instead running from top to bottom (direction illustrated by red arrows). Thick blue lines indicate areas of thickening. Image reconstructions occurred in Drishti *(a)* and ORS Dragonfly *(b*). Segmentation of pores occurred in Zeiss Microscopy Intellesis software. Pore thickness map produced by Local Thickness plugin in Fiji/ImageJ software.

Segmentation of the pores using Intellesis machine learning segmentation software (Zeiss Microscopy, Oberkocken, Germany) reveals the morphology of pores in 3D through the length of the ala. Nearest the apex the pores form fan-like networks which continues down into the ala length (figure 5 *b i, ii*). However towards the ala sutural edge and base the morphology and orientation of the pores change, and are instead parallel to the ala surface running from top to bottom; a simplified diagram of this is seen in (figure 5 *b iv*). In addition, there is a widening of pores on the inside edge of the plate nearest the soft bodied organism (figure 5 *b iii*). Despite the identification of pores, no grain boundaries, crystal structure or segregated grayscale variations were observed via XRM imaging, therefore requiring further characterisation via other techniques (e.g., FIB-SEM, EBSD, SEM).

### 4.3. Pore nano-structure (FIB-SEM) and quantification

From targeted FIB-SEM nano-tomographic milling through Atlas 5/3D (*section 3.4.1*), it is possible to study the ala pore networks at a higher resolution to establish nano-scale features and relationships. We have compared ala pore networks with those on the opposing plate (figure 6) to establish exoskeleton variations in pore structure. 10 × 10 µm FIB-SEM nano-tomographic volumes reveal variations in pore morphology and alignment between those on the ala and those on the neighbouring plate (figure 6). Pores on the ala (figure 6 *d, e*) are numerous (962 in this volume), have pore diameters up to 1.56µm, and are composed of mostly shorter and singular pores. This is compared with those on the opposing plate (figure 6 *b, c*) which are less numerous (594 in this volume) and are dominated by thicker and longer connected pores. Ala pore directionality (figure 6 *d, f*) follows EBSD crystallographic orientations (figure 4 *e, f*), however dominant orientations on the opposing plate (figure 6 *b, c*) do not appear to be related to crystallographic structure (figure 4 *e, f*). Further analysis to the porosities via Avizo software indicates similar trends in elongation and pore diameter between the opposing plate and the ala (figures 6 *f, i*), with a larger spread for values of pore width (figure 6 *h*) and more spherical pores (figure 6 *g*) in the ala. This illustrates that individual pores and pore networks vary in structure (and possibly function) across the barnacle shell. No crystal structure, crystal boundaries or phase variations were observed from FIB-SEM images.

**Figure 6:**
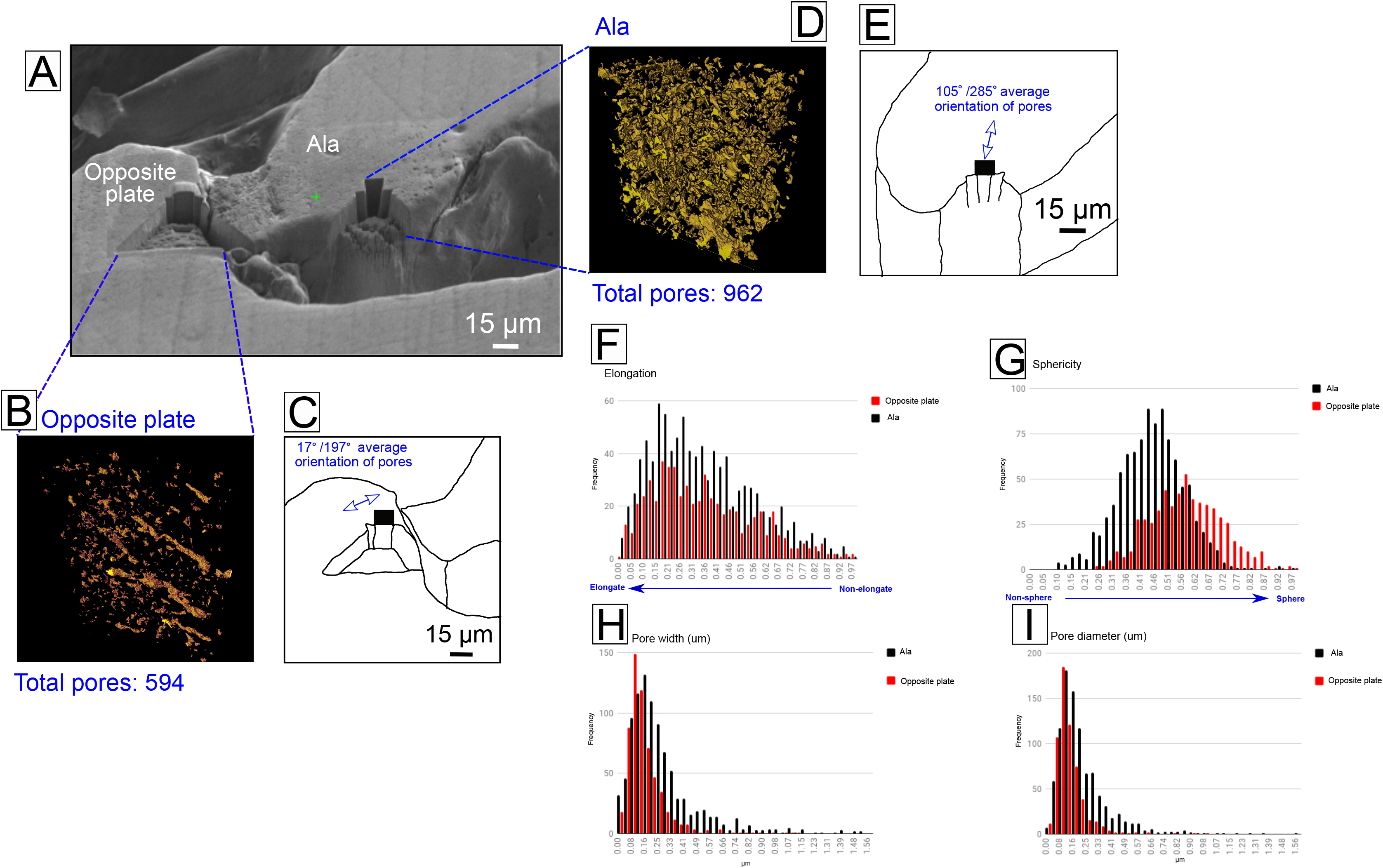
*(a)* Locations of milled volumes on the ala and opposing plate. (*b, c*) Reconstructed and segmented pore volumes on the opposing plate, illustrating a 17/197° orientation. (*d, e*) Reconstructed and segmented pore volumes on the ala, illustrating a 105/285° orientation. *(f-i)* Histograms highlighting statistical variations in the pores between the ala and opposing plate.

## 5. Discussion

### 5.1. Correlating multi-modal and multi-scale data/images

This work represents the first correlative multi-modal and multi-scale study of barnacle morphological and mechanical structure across multiple dimensions. Correlative microscopy overcomes the multi-scale ‘needle in a haystack’ challenge of working in complex 3D volumes, and has proved successful for accurately locating specific regions of study in human-made materials; examples include lithium ion batteries [35] and corrosion in magnesium alloys [36]. Additionally, the technique is well established across a range of applications in the life sciences [37–40]. The advantage of using a multi-modal and correlative approach is that each specialised technique can provide information relating to a specific feature or structure, and that correlation across dimensions can thus inform how features and structures are linked, particularly in hierarchical materials. Increasing the resolution is important for identifying and improving the accuracy of measurement of features at the micro to nano-scale (e.g. the voids in figure 5 *b*, 6) and in three dimensions reveals characteristics that might not be identifiable in one or two dimensions alone (e.g. pore orientations in figure 5).

The correlative workflow improves our understanding of barnacle shell structure (figures 6, 7) where specific regions are accurately located to the nano-scale. The workflow outlined here can be utilised in other bioinspiration studies (e.g., mollusc shells) to correlate macro to nano-scale shell structures, which ultimately improves the understanding of form and function as well as application for human-made materials.

**Figure 7:**
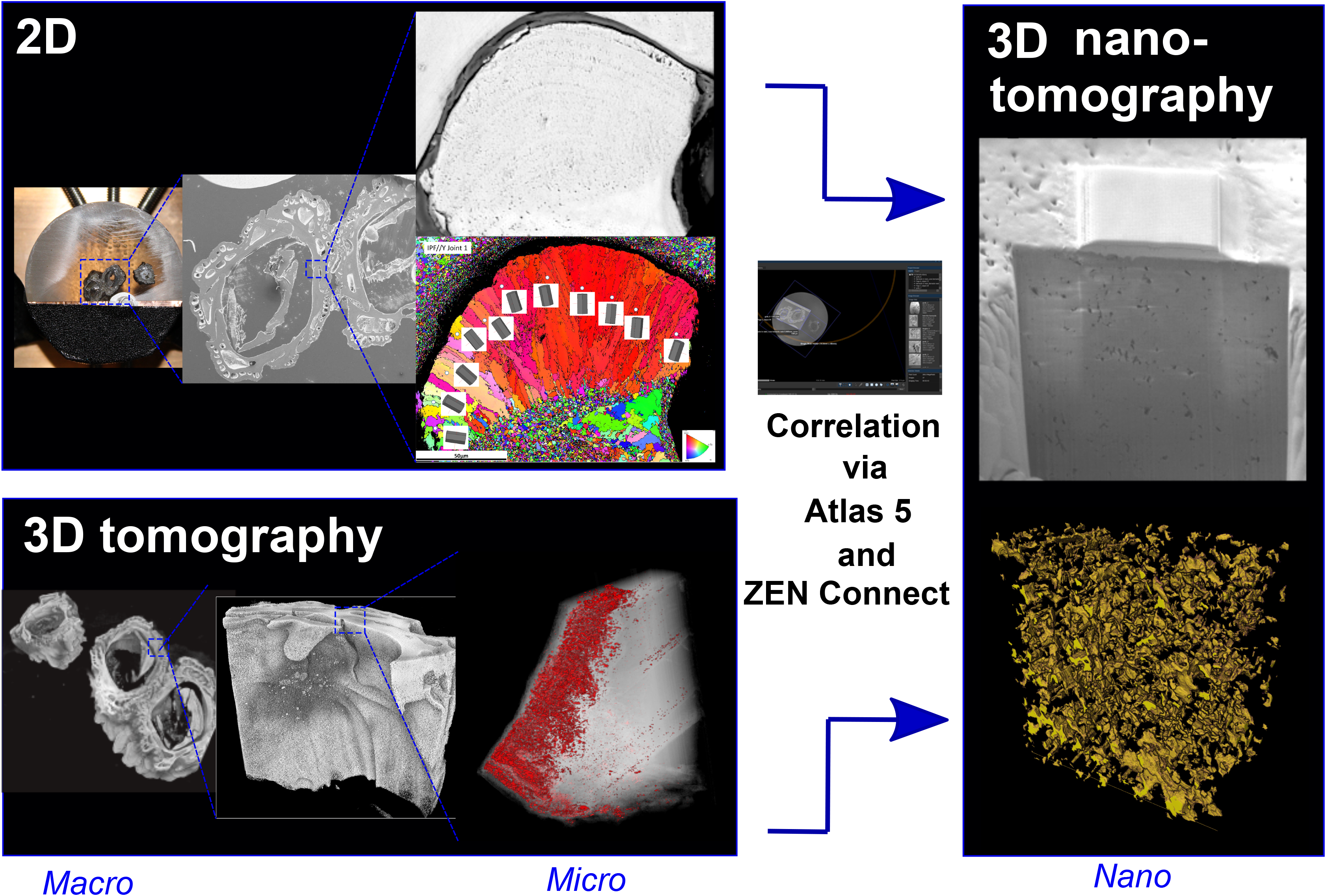
Correlation of 2D and 3D over macro-micro-nano scales and multi-modes to inform about barnacle exoskeleton morphology.

### 5.2. Crystallographic alignment with pores

This work reveals that the ala tips of *S. balanoides* exhibit a distinct crystallographically-graded biological material (figure 4). We propose the elongated crystal growth at the tip of the ala compared to the more equi-axed grains behind the tip and on the opposing plate (Figures 4 *e, f*) represent a growing front in a region of active biomineralisation. During biomineralisation, calcium carbonate generally forms prismatic, sheet nacreous, lenticular nacreous, foliated, cross-lamellar, and homogeneous crystal morphologies [41]. Only prismatic and homogeneous crystals were identified here in barnacles (figure 4 e, f). Elongated crystals in *Semibalanus* alae have previously been identified from a single study of *B. balanus* and *S. balanoides* [17] that was limited to two dimensional study (LM and SEM) and left their origins unexplained, and elongate crystals have been identified in *B. amphitrite* [19,21]. Different crystallographic orientations, in particular elongate, prismatic columns associated with organic materials are common in a variety of biomineralised molluscs [6,42,43] but remain largely unidentified in barnacles. The ordering of calcite in the scutum (one of the two plates that guard the apical opercular opening) is significantly disordered compared with the calcite in the wall-plates *in A. amphitrite* [19], and the calcitic microcrystals in the wall plates of this species show almost no orientation [19,21] whilst those in the base plate of *A. amphitrite* shows some preferred alignment [21]. This suggests there may be some variations between barnacle genera other than *Semibalanus balanoides*. Elongated, prismatic calcite columns growing perpendicularly to the shell surface are present in shells of various molluscs (including oysters and scallops; [42,44]) and other arthropods (specifically the Mantis shrimp; [45]), which are surrounded by up to 3 µm-thick organic membranes and vaterite columns in the shell of the bivalve *Corbicula fluminea* [6]. This indicates that different crystallographic orientations, in particular elongate, prismatic columns associated with organic materials are common in a variety of biomineralised molluscs, however remain largely unexplained and undescribed in barnacles and may form an important part of the shell structure. Several hypotheses, either independently or in union, may explain the crystallographic elongation at the tip of the ala in *S. balanoides*:

1. Elongation could be related to the calcium carbonate polymorph that is being biomineralised (e.g., aragonite/vaterite/calcite) which may form specific morphologies [46]. Calcite is the most stable polymorph, with aragonite forming at high pressures and vaterite being thermodynamically unstable [46,47]. Extant barnacle shells are all reportedly dominated by calcite [4], but were all originally phosphatic [48,49]; only one extant species now uses calcium phosphate (*Ibla cumingi;* [50]) but a detailed study of variations within exoskeletons and between genera/species has never been undertaken. Changes in the form of calcium carbonate/calcium phosphate could impact the mechanical structure and integrity of different areas of the exoskeleton, and of barnacles of different chemistries/polymorphs. Some molluscs biomineralise aragonite instead of calcite in seawater rich in magnesium [46,51], so the specific habitat/latitude of different barnacles could also affect crystal structure and mechanical properties of the shell.
2. The age/growth stage of the organism. The transverse banding forming perpendicularly to the elongate crystal orientation (figure 4 *c*, *d*, 5 *a*) is postulated by [17] to indicate variations in growth rate (like tree rings). Acorn barnacles such as *S. balanoides* grow by lengthening their side (wall) plates [20] and biomineralising the base of their exoskeleton in an incremental fashion [52]. Therefore, the transverse banding at the ala tip probably indicates incremental growth spurts and the perpendicular elongate crystals are younger than those biomineralised as smaller, more equiaxed grains (figure 4 *e*, *f*). [20] suggests crystallisation initiates at the leading edge of the barnacle base plate with the deposition of mutually aligned fine-grained calcite, which then acts as a template for the formation of subsequent coarser grains. A similar process may occur in the wall plates and alae of *S. balanoides*, with elongate crystals growing upon finer-grained granular calcite (figure 4 *e, f*). *S. balanoides* shows little growth after 1-2 years [17], however in our specimens it is unclear whether the barnacle was still growing or fully formed. Also unclear is whether the elongate ‘growth’ crystals are overprinted later in life. Comparably, molluscs biomineralise their shells continuously whereas barnacles do not [53], indicating that crystal growth in the barnacle occurs much quicker than molluscs, possibly leading to unique crystallographic configuration morphologies.
3. Even though we do not see an obvious organic layer separating the tip of the ala from the neighbouring plate in this study (figure 4 *c, d*). [25] state that an organic layer between the two plates enables them to ‘stick’ together. This could be an important feature as organic material can promote biomineralization, crystallographic morphology and orientation, and ultimately contribute towards exoskeleton mechanical properties [4,42,54,55]. Biomineralised structures are not purely inorganic because they all contain some organic molecules within their structure [42] and hydrogels often provide biological control on the construction of aligned calcium carbonate domains [19,20]. In many marine shell-producing organisms, the hydrogel slows grain motion enabling carbonate grains to orientate themselves relative to each other [20,47]. Indeed, the crystal properties and microstructure in *A. amphitrite* are consistent with those developing in a hydrogel-like environment and the intercrystalline organic matrix is a non-proteinaceous sulphate-rich polymer behaving like a hydrogel [19]. An organic matrix is presumed responsible for the organisation of exoskeleton biomineralisation in the giant barnacle *Austromegabalanus psittacus* where it controls the type, size and orientation of exoskeleton-forming crystals [53]. Consequently, it could be inferred that organic matrices have an influential effect on biomineralisation in barnacles and might affect crystal shape and size, and through this the mechanical properties of the exoskeleton.

### 5.3. Pore networks represent organic channels and ‘pockets’?

We have identified and examined numerous porous channels in the barnacle alae. We considered ala pores may represent crystallographic boundaries (figure 4 *e, f*) as they have the same orientation (figure 6, *d, e*) however further study via FIB-SEM showed the pores sit within the grains (some crystals are up to 10µm wide, the same as the entire FIB-SEM volumes; figure 4 *e, f*, 6 *d*). This could be a factor of the EBSD resolution, however it is likely the pores exist independently of crystallographic structure whilst maintaining the same orientations. This hypothesis is supported by elongate pore networks in the more equi-axed crystals of the opposing plate and behind the ala tip (figure 6 *b*) and the lack of observed crystallography in the cross-section face during FIB-SEM milling (figure 5 *c*), suggesting that the locations for FIB-SEM nanotomographic milling were small enough to be considered intragrain.

Exoskeleton/shell pores are common in many groups of biomineralised marine organisms including gastropods [56], bivalves [6,57], and within the exoskeleton of some barnacles (particularly in base plates; [25,50]). It has been suggested that the most important adaptive breakthrough in balanoid barnacles, and their competitive success over *Chthamalus* barnacles, is a tubiferous wall structure which enables fast exoskeletal growth and colonisation of free space [7]. We propose that the ala pores in the exoskeleton of *B. balanoides* are organic-rich areas, possibly involved in biomineralisation. The involvement of organic material in the biomineralisation of specific crystal structures and orientations could have a bearing on the function of the ala pores, and may represent channels/canals which hold or deliver biomineralisation products to specific areas of the exoskeleton. Organic membranes are known to influence the pattern of columnar prismatic layers in numerous mollusc shells [43], so it is possible that organic channels (or, ala pores) running through the barnacle structure contribute towards the delivery of and biomineralisation of calcium carbonate. The organic layer separating the ala tip and neighbouring plate may play a part in this. A layer of organic cuticle exists between the ala and neighbouring plate in *Balanus balanus* [17] which may explain the concentration of pores and elongated crystals near the ala tip. The pores, however, are not all elongate channels and some pores, particularly in the opposing plate, being more spherical in shape (figure 6 *g*). Pores in different parts of the exoskeleton may therefore have different functions, possibly acting as channels in the ala to deliver organic material for biomineralisation, and to hold pockets of organic material in the opposing plate.

Barnacle wall-plate mineralisation occurs through cell-mediated Ca^2+^ uptake, storage and mobilization to the mineralization front [32] and pore canals assist in transporting components necessary for calcification (including Ca^2+^ and organic molecules). Voids/pores in the aragonitic platelets of mollusc nacre contain increased amounts of carbon [58] and tube-like shell pores containing organic material are also present in limpets [57] whilst the organic intertile layer in abalone is anchored by the growth of minerals through pores [59]. The pores forming ‘canal’ networks in the wall plates of large sessile barnacles (*Austromegabalanus psittacus;* [24]) has yet to be ascribed a function. Longitudinal canals in the wall plates, and radial canals in the base plate of *B. amphitrite* are lined with mantle epithelium, and biomineralisation is facilitated by salt-rich secretions from the junction between the wall and base plates [32]. Some barnacles also possess microducts/pores in their base plates to facilitate secretion of adhesive [25]. Exoskeleton pores seem to be used for the transport of organic material and biomineralisation, although the role of proteins and other macromolecules in the biomineralisation process however is still poorly understood [19], and future studies should aim to quantify this.

### 5.4. Variation in shell morphology in barnacles

Despite the presence of probable organic pore networks and specific crystallographic orientations in many genera and species of barnacle [17,24,25,29], it is possible the features discovered in this study are unique to *S. balanoides*. The alae and wall plates of other barnacles, for example *B. balanus*, are considerably different to those of *S. balanoides* [17], consequently their crystallographic structures may also differ. Shell morphology is also highly phenotypically plastic within a barnacle species and can change according to wave exposure [60], predation [61], and, especially, crowding [62]. Hummocks of tall, columnar barnacles are common under high population densities whilst squat, conical growth forms with much thicker (2-5 times in *S. balanoides*) walls dominate in solitary individuals or low population densities [62]. Whether these growth forms differ in microstructural and mechanical properties may warrant investigation, although the substantial difference in shell strength between *B. balanus* and *S. balanoides* has been attributed to distinct variation in shell architecture rather than mechanical properties of the wall plates [17].

### 5.5. Implications for mechanical strength and bioinspiration

The range of crystal sizes and shapes, as well as reinforcement by organic-rich channels, will all contribute to the mechanical properties of the barnacle shell. For example, the probable organic pores and specifically oriented crystallographic structure of the ala tip may also act as a strengthening mechanism in a region of active growth [20] and high stress [60,61]. The presence of organic material within the biomineralised structure also has important implications for strength and toughness. Crossed-lamellar structures, composed of aragonite and a small amount of organic material, are the most common microstructures in mollusc shells and possess a fracture toughness and elasticity much higher than pure carbonate (calcite) mineral [58,63–66]. Indeed, removal of this organic material from abalone shell greatly contributes towards its mechanical decline [67]. The ala in *S. balanoides* is non-geometric through its length (figures 1, 5 *a-b*; [17]) suggesting it is potentially not the strongest design for an interlocking joint. Further tests are required to establish the hardness and strength of different regions of the barnacle, in particular, the alae, and the effect the elongated crystal structure and organic-rich pores of specific orientations have on mechanical strength.

Understanding the morphology and structure of biomaterials can contribute towards the design and manufacture of human-made materials [2,68,69]. Similar discrete bimodal grain sizes are observed in manufactured materials for aerospace, such as the ‘dual microstructure’ of some nickel superalloy-based gas turbine disks [70]. To improve the material properties and in-service behaviour, the material is specifically designed to have distinct microstructures in different regions of the disk. A fine-grained microstructure is produced in the bore, providing a higher proof strength and fatigue life; whereas a coarse-grained microstructure in the rim results in greater creep life [71]. Location-specific microstructures in different regions of the part are tuned to the environmental conditions in which they are exposed for optimised design and life. But this level of modification to different parts of the microstructure requires multiple complex steps, including heat treatments at temperatures in excess of 650ºC [72]. It is possible the barnacle alae dual microstructure with specific crystallographic orientation of the elongated grains perpendicular to the loading/contact surface is a functional characteristic, with highly adapted microstructural features driven by evolutionary processes. In comparison to the processes required to produce the nickel superalloy, the barnacle achieves a highly-complex microstructure in ambient conditions, dynamic tidal conditions, and with the chemistry and temperatures imposed on it by the environment.

Additionally, the interlocking nature of the barnacle joints described here, combined with the variation in crystallographic organisation and pore structure, could contribute towards the development of materials that require movement and expansion whilst remaining strong, such as expandable pressurised containers or submersible structures. Another potential could be the utilisation of barnacle-like designs in additive manufacturing. In recent years Functionally Graded Additive Manufacturing (FGAM) has developed its capabilities of fabricating materials layer by layer and by controlling morphological features (such as porosity) to create structurally and mechanically distinctive materials [73]. A correlative approach to understanding the morphological, chemical, and structural characteristics of natural biomaterials outlined in this study could therefore contribute greatly to the development of future bioinspired materials.

## 6. Conclusions

Here we show the advantages of using multi-modal, multi-dimensional and multi-scale correlative microscopy techniques to identify the morphological, microstructural, and crystallographic properties of the shell of the barnacle *S. balanoides*. The barnacle shell is composed of six interlocking calcium carbonate wall-plates with alae (*supplementary material 2)*, finger-like protrusions acting as a contact point of potential high stress for the joining of adjacent plates. 2D imaging via LM and SEM indicate that the tip of the ala contains a series of pores, and from EBSD we illustrate the crystallographically-graded texture of the biomineralised calcium carbonate, with the elongated grains at the outer edge of the ala oriented perpendicularly to the contact surface of the joint, and the c-axis rotated with the radius of the ala; the same orientation as the pores. 3D imaging via XRM enables the segmentation of the pores, and the realisation that pores are only visible within the very tip of the ala, their orientations change through its length, and there is pore thickening on the inside (soft bodied) edge of the plate. Further analysis of the nano-scale structure of the pores through FIB-SEM illustrate that the pores are probably organic channels and pockets which are involved with the biomineralisation process. These properties indicate the macro-micro-nano scale features of the barnacle exoskeleton, particularly at the ala, could be useful for bioinspiration for human-made materials. Furthermore, correlative imaging allows the targeting of specific regions of interest across different imaging techniques and length scales, and greatly increases the amount of information that can be acquired from imaging in purely two dimensions, bridging the materials science of structure-property relationships with the biological form and function.

## Supporting information

Supplemental 1 - Avizo porosity data

Supplemental 2 - Barnacle plate video

Supplemental 3 - Ala video

Supplemental 4 - Opposite joint video

## 7. Acknowledgements

We thank two anonymous reviewers for insightful reviews and comments on this manuscript. Additional thanks go to James Russell and Imogen Woodhead for aid during SEM imaging and preliminary pore analysis respectively, both from Swansea University, and Tobias Volkenandt and Stefanie Freitag from Carl Zeiss Microscopy (Germany).

## 9. Data accessibility

Supplementary material (1-4) can be found at X. XRM scans (tiff stacks) of whole barnacles mounted in resin and individual plate can be found at X (will be Dryad repository).

**Supplementary material 1:** Data for ala porosity analysis, generated from FIB-SEM imaging.

**Supplementary material 2:** Video illustrating the 3D structure of the barnacle and the adjoining wall-plates, generated from XRM. Rendered in Drishti.

**Supplementary material 3:** Video illustrating the pore networks in the ala; data generating from FIB-SEM imaging. Rendered in ORS Dragonfly.

**Supplementary material 4:** Video illustrating the pore networks in the opposing plate to the ala; data generating from FIB-SEM imaging. Rendered in ORS Dragonfly.

## References

1. Albéric M, Bertinetti L, Zou Z, Fratzl P, Habraken W, Politi Y. 2018 The Crystallization of Amorphous Calcium Carbonate is Kinetically Governed by Ion Impurities and Water. Adv. Sci. 5. (doi:10.1002/advs.201701000)

2. North L, Labonte D, Oyen ML, Coleman MP, Caliskan HB, Johnston RE. 2017 Interrelated chemical-microstructural-nanomechanical variations in the structural units of the cuttlebone of *Sepia officinalis*. APL Mater. 5, 116103. (doi:10.1063/1.4993202)

3. Gal A, Kahil K, Vidavsky N, Devol RT, Gilbert PUPA, Fratzl P, Weiner S, Addadi L. 2014 Particle accretion mechanism underlies biological crystal growth from an amorphous precursor phase. Adv. Funct. Mater. 24, 5420–5426. (doi:10.1002/adfm.201400676)

4. Astachov L, Nevo Z, Brosh T, Vago R. 2011 The structural, compositional and mechanical features of the calcite shell of the barnacle Tetraclita rufotincta. J. Struct. Biol. 175, 311–318. (doi:10.1016/j.jsb.2011.04.014)

5. Ma Y et al. 2009 The grinding tip of the sea urchin tooth exhibits exquisite control over calcite crystal orientation and Mg distribution. Proc. Natl. Acad. Sci. 106, 6048–6053. (doi:10.1073/pnas.0810300106)

6. Frenzel M, Harrison RJ, Harper EM. 2012 Nanostructure and crystallography of aberrant columnar vaterite in *Corbicula fluminea* (Mollusca). J. Struct. Biol. 178, 8–18. (doi:10.1016/j.jsb.2012.02.005)

7. Stanley SM, Newman WA. 1980 Competitive exclusion in evolutionary time: the case of the acorn barnacles. Paleobiology 6, 173–183.

8. Tomanek L, Helmuth B. 2002 Physiological Ecology of Rocky Intertidal Organisms: A Synergy of Concepts. Integr. Comp. Biol. 42, 771–775. (doi:10.1093/icb/42.4.771)

9. Burden DK, Spillmann CM, Everett RK, Barlow DE, Orihuela B, Deschamps JR, Fears KP, Rittschof D, Wahl KJ. 2014 Growth and development of the barnacle *Amphibalanus amphitrite□*: time and spatially resolved structure and chemistry of the base plate. Biofouling 30, 799–812. (doi:10.1080/08927014.2014.930736)

10. R.M. O’Riordan, Power AM, Myers AA. 2010 Factors, at different scales, affecting the distribution of species of the genus *Chthamalus ranzani* (Cirripedia, Balanomorpha, Chthamaloidea). J. Exp. Mar. Bio. Ecol. 392, 46–64.

11. du Plessis A, Broeckhoven C. 2019 Looking deep into nature: A review of micro-computed tomography in biomimicry. Acta Biomater. 85, 27–40. (doi:10.1016/j.actbio.2018.12.014)

12. Nakamura K, Hisanaga T, Fujimoto K, Nakajima K, Wada H. 2018 Plant-inspired pipettes. J. R. Soc. Interface 15, 20170868. (doi:10.1098/rsif.2017.0868)

13. Cutkosky MR. 2015 Climbing with adhesion: From bioinspiration to biounderstanding. Interface Focus 5, 20150015. (doi:10.1098/rsfs.2015.0015)

14. Porter MM, Imperio R, Wen M, Meyers MA, McKittrick J. 2014 Bioinspired scaffolds with varying pore architectures and mechanical properties. Adv. Funct. Mater. 24, 1978–1987. (doi:10.1002/adfm.201302958)

15. Barthelat F. 2007 Biomimetics for next generation materials. Philos. Trans. R. Soc. A Math. Phys. Eng. Sci. 365, 2907–2919. (doi:10.1098/rsta.2007.0006)

16. Ripley RL, Bhushan B. 2016 Bioarchitecture: Bioinspired art and architecture-a perspective. (doi:10.1098/rsta.2016.0192)

17. Murdock GR, Currey JD. 1978 Strength and design of shells of the two ecologically distinct barnacles, *Balanus balanus* and *Semibalanus (Balanus) balanoides* (Cirripedia). Biol. Bull. 155, 169–192.

18. Clare AS, Høeg JT. 2008 *Balanus amphitrite* or *Amphibalanus amphitrite?* A note on barnacle nomenclature. Biofouling 24, 55–57. (doi:10.1080/08927010701830194)

19. Khalifa GM, Weiner S, Addadi L. 2011 Mineral and matrix components of the operculum and shell of the barnacle *Balanus amphitrite*: Calcite crystal growth in a hydrogel. Cryst. Growth Des. 11, 5122–5130. (doi:10.1021/cg2010216)

20. De Gregorio BT, Stroud RM, Burden DK, Fears KP, Everett RK, Wahl KJ. 2015 Shell Structure and Growth in the Base Plate of the Barnacle *Amphibalanus amphitrite*. ACS Biomater. Sci. Eng. 1, 1085–1095. (doi:10.1021/acsbiomaterials.5b00191)

21. Lewis AC, Burden DK, Wahl KJ, Everett RK. 2014 Electron Backscatter Diffraction (EBSD) study of the structure and crystallography of the barnacle *Balanus amphitrite*. J. Miner. Met. Mater. Soc. 66, 143–148. (doi:10.1007/s11837-013-0793-y)

22. Sangeetha R, Kumar R, Venkatesan R, Doble M, Vedaprakash L, Kruparatnam, Lakshmi K, Dineshram. 2010 Understanding the structure of the adhesive plaque of *Amphibalanus reticulatus*. Mater. Sci. Eng. C 30, 112–119. (doi:10.1016/j.msec.2009.09.007)

23. Hui CY, Long R, Wahl KJ, Everett RK. 2011 Barnacles resist removal by crack trapping. J. R. Soc. Interface 8, 868–879. (doi:10.1098/rsif.2010.0567)

24. Fernandez MS, Vergara I, Oyarzum A, Arias JI, Rodriguez R, Wiff JP, Fuenzalida VM, Arias JL. 2002 Extracellular Matrix Molecules Involved in Barnacle Shell Mineralization. Mater. Res. Soc. Symp. Proc. 724, 1–9.

25. Raman S, Kumar R. 2011 Construction and nanomechanical properties of the exoskeleton of the barnacle, *Amphibalanus reticulatus*. J. Struct. Biol. 176, 360–369. (doi:10.1016/j.jsb.2011.08.015)

26. Slater TJA, Bradley RS, Bertali G, Geurts R, Northover SM, Burke MG, Haigh SJ, Burnett TL, Withers PJ. 2017 Multiscale correlative tomography: An investigation of creep cavitation in 316 stainless steel. Sci. Rep. 7, 1–10. (doi:10.1038/s41598-017-06976-5)

27. Daly M, Burnett TL, Pickering EJ, Tuck OCG, Léonard F, Kelley R, Withers PJ, Sherry AH. 2017 A multi-scale correlative investigation of ductile fracture. Acta Mater. 130, 56–68. (doi:10.1016/j.actamat.2017.03.028)

28. Grandfield K, Engqvist H. 2012 Focused ion beam in the study of biomaterials and biological matter. Adv. Mater. Sci. Eng. 2012, 1–6. (doi:10.1155/2012/841961)

29. Mitchell RL, Pleydell-Pearce C, Coleman MP, Davies P, North L, Johnston RE, Harris W. 2018 Correlative Imaging and Bio-inspiration: Multi-scale and Multi-modal Investigations of the Acorn Barnacle *(Semibalanus balanoides*). Microsc. Microanal. 24, 376–377. (doi:10.1017/s1431927618002374)

30. Mitchell R., Coleman M, Davies P, North L, Pope E., Pleydell-Pearce C, Harris W, Johnston R. 2019 Macro-to-nano scale investigation of the acorn barnacle *Semibalanus balanoides*: correlative imaging, biological form and function, and bioinspiration. In Biorxiv,

31. Hayward PJ, Ryland JS. 2017 Handbook of the marine fauna of north-west Europe (2nd edition). 2nd edn. Oxford University Press, Oxford.

32. Gohad N V., Dickinson GH, Orihuela B, Rittschof D, Mount AS. 2009 Visualization of putative ion-transporting epithelia in *Amphibalanus amphitrite* using correlative microscopy: Potential function in osmoregulation and biomineralization. J. Exp. Mar. Bio. Ecol. 380, 88–98. (doi:10.1016/j.jembe.2009.09.008)

33. Anderson DT. 1994 Barnacles: Structure, Function, Development and Evolution. Chapman and Hall, London.

34. Pilati T, Demartin F, Gramaccioli CM. 1998 Lattice estimation of atomic displacement parameters in carbonates: calcite and aragonite CaCO3, dolomite (CaMg(CO3)2 and magnesite MgCO3. Acta Cryst. B54, 515–523.

35. Gelb J, Finegan DP, Brett DJL, Shearing PR. 2017 Multi-scale 3D investigations of a commercial 18650 Li-ion battery with correlative electron- and X-ray microscopy. J. Power Sources 357, 77–86. (doi:10.1016/j.jpowsour.2017.04.102)

36. Harris W. 2015 Multi-scale Correlative Study of Corrosion Evolution in a Magnesium Alloy.

37. Bradley RS, Withers PJ. 2016 Correlative multiscale tomography of biological materials. MRS Bull. 41, 549–556. (doi:10.1557/mrs.2016.137)

38. Sykes D, Hartwell R, Bradley RS, Burnett TL, Hornberger B, Garwood RJ, Withers PJ. 2019 Time-lapse three-dimensional imaging of crack propagation in beetle cuticle. Acta Biomater. 86, 109–116. (doi:10.1016/j.actbio.2019.01.031)

39. Bernhardt M et al. 2018 Correlative microscopy approach for biology using X-ray holography, X-ray scanning diffraction and STED microscopy. Nat. Commun. 9, 1–9. (doi:10.1038/s41467-018-05885-z)

40. Handschuh S, Baeumler N, Schwaha T, Ruthensteiner B. 2013 A correlative approach for combining microCT, light and transmission electron microscopy in a single 3D scenario. Front. Zool. 10, 1–16. (doi:10.1186/1742-9994-10-44)

41. Liang Y, Zhao Q, Li X, Zhang Z, Ren L. 2016 Study of the microstructure and mechanical properties of white clam shell. Micron 87, 10–17. (doi:10.1016/j.micron.2016.04.007)

42. Okumura T, Suzuki M, Nagasawa H, Kogure T. 2010 Characteristics of biogenic calcite in the prismatic layer of a pearl oyster, *Pinctada fucata*. Micron 41, 821–826. (doi:10.1016/j.micron.2010.05.004)

43. Checa AG, Macías-Sánchez E, Harper EM, Cartwright JHE. 2016 Organic membranes determine the pattern of the columnar prismatic layer of mollusc shells. Proc. R. Soc. B Biol. Sci. 283. (doi:10.1098/rspb.2016.0032)

44. Rodriguez-Navarro AB, Checa A, Willinger M-G, Bolmaro R, Bonarski J. 2012 Crystallographic relationships in the crossed lamellar microstructure of the shell of the gastropod *Conus marmoreus*. Acta Biomater. 8, 830–835. (doi:10.1016/j.actbio.2011.11.001)

45. Weaver JC et al. 2012 The Stomatopod Dactyl Club: A formidable damage-tolerant biological hammer. Science 336, 1275–1280. (doi:10.1126/science.1218764)

46. Falini G, Albeck S, Weiner S, Addadi L. 1996 Control of Aragonite or Calcite Polymorphism by Mollusk Shell Macromolecules. Science. 271, 67–69.

47. Zhou GT, Yao QZ, Ni J, Jin G. 2009 Formation of aragonite mesocrystals and implication for biomineralization. Am. Mineral. 94, 293–302. (doi:10.2138/am.2009.2957)

48. Gale A. 2018 Stalked barnacles (Cirripedia, Thoracica) from the Upper Jurassic (Tithonian) Kimmeridge Clay of Dorset, UK; palaeoecology and bearing on the evolution of living forms. Proc. Geol. Assoc., 1–11. (doi:10.1016/j.pgeola.2018.01.005)

49. Pérez-Losada M, Harp M, Høeg JT, Achituv Y, Jones D, Watanabe H, Crandall KA. 2008 The tempo and mode of barnacle evolution. Mol. Phylogenet. Evol. 46, 328–346. (doi:10.1016/j.ympev.2007.10.004)

50. Lowenstam HA, Weiner S. 1992 Phosphatic shell plate of the barnacle *Ibla* (Cirripedia): a bone-like structure. Proc. Natl. Acad. Sci. 89, 10573–10577. (doi:10.1073/pnas.89.22.10573)

51. Checa AG, Jiménez-López C, Rodríguez-Navarro A, Machado JP. 2007 Precipitation of aragonite by calcitic bivalves in Mg-enriched marine waters. Mar. Biol. 150, 819–827. (doi:10.1007/s00227-006-0411-4)

52. Hockett D, Ingram P, LeFurgey A. 1997 Strontium and manganese uptake in the barnacle shell: Electron probe microanalysis imaging to attain fine temporal resolution of biomineralization activity. Mar. Environ. Res. 43, 131–143. (doi:10.1016/0141-1136(96)00081-5)

53. Rodriguez-Navarro AB, CabraldeMelo C, Batista N, Morimoto N, Alvarez-Lloret P, Ortega-Huertas M, Fuenzalida VM, Arias JI, Wiff JP. 2006 Microstructure and crystallographic-texture of giant barnacle (*Austromegabalanus psittacus*) shell. J. Struct. Biol. 156, 355–362. (doi:10.1016/j.jsb.2006.04.009)

54. Arias JL, Neira-Carrillo A, Arias JI, Escobar C, Bodero M, David M, Fernández MS. 2004 Sulfated polymers in biological mineralization: A plausible source for bio-inspired engineering. J. Mater. Chem. 14, 2154–2160. (doi:10.1039/b401396d)

55. Bezares J, Asaro RJ, Hawley M. 2010 Macromolecular structure of the organic framework of nacre in *Haliotis rufescens*: Implications for mechanical response. J. Struct. Biol. 170, 484–500. (doi:10.1016/j.jsb.2010.01.006)

56. Heß M, Beck F, Gensler H, Kano Y, Kiel S, Haszprunar G. 2008 Microanatomy, shell structre and molecular phylogeny of *Leptogyra, Xyleptogyra* and *Leptogyropsis* (Gastropoda: Neomphalida: Melanodrymiidae) from sunken wood. J. Molluscan Stud. 74, 383–401. (doi:10.1093/mollus/eyn030)

57. Reindl S, Haszprunar G. 1994 Light and electron microscopical investigations on shell pores (caeca) of fissurellid limpets (Mollusca: Archaeogastropoda). J. Zool. Soc. London 233, 385–404. (doi:10.1111/j.1469-7998.1994.tb05272.x)

58. Gries K, Kröger R, Kübel C, Fritz M, Rosenauer A. 2009 Investigations of voids in the aragonite platelets of nacre. Acta Biomater. 5, 3038–3044. (doi:10.1016/j.actbio.2009.04.017)

59. Meyers MA, Lim CT, Li A, Hairul Nizam BR, Tan EPS, Seki Y, McKittrick J. 2009 The role of organic intertile layer in abalone nacre. Mater. Sci. Eng. C 29, 2398–2410. (doi:10.1016/j.msec.2009.07.005)

60. Pentcheff ND. 1991 Resistance to crushing from wave-borne debris in the barnacle Balanus glandula. Mar. Biol. 110, 399–408. (doi:10.1007/BF01344359)

61. Lively CM. 2006 Predator-Induced Shell Dimorphism in the Acorn Barnacle *Chthamalus anisopoma*. Evolution 40, 232–242. (doi:10.2307/2408804)

62. Bertness MD, Gaines SD, Yeh SM. 1998 Making mountains out of barnacles: The dynamics of acorn barnacle hummocking. Ecology 79, 1382–1394. (doi:DOI: 10.2307/176750)

63. Quan H, Yang W, Schaible E, Ritchie RO, Meyers MA. 2018 Novel Defense Mechanisms in the Armor of the Scales of the “Living Fossil” Coelacanth Fish. Adv. Funct. Mater. 28, 1804237. (doi:10.1002/adfm.201804237)

64. Li XW, Ji HM, Yang W, Zhang GP, Chen DL. 2017 Mechanical properties of crossed-lamellar structures in biological shells: A review. J. Mech. Behav. Biomed. Mater. 74, 54–71. (doi:10.1016/j.jmbbm.2017.05.022)

65. Liu Z, Meyers MA, Zhang Z, Ritchie RO. 2017 Functional gradients and heterogeneities in biological materials: Design principles, functions, and bioinspired applications. Prog. Mater. Sci. 88, 467–498. (doi:10.1016/j.pmatsci.2017.04.013)

66. Liu Z, Zhu Y, Jiao D, Weng Z, Zhang Z, Ritchie RO. 2016 Enhanced protective role in materials with gradient structural orientations: Lessons from Nature. Acta Biomater. 44, 31–40. (doi:10.1016/j.actbio.2016.08.005)

67. Lopez MI, Meza Martinez PE, Meyers MA. 2014 Organic interlamellar layers, mesolayers and mineral nanobridges: Contribution to strength in abalone (*Haliotis rufescence*) nacre. Acta Biomater. 10, 2056–2064. (doi:10.1016/j.actbio.2013.12.016)

68. Wilkerson RP, Gludovatz B, Ell J, Watts J, Hilmas GE, Ritchie RO. 2019 High-temperature damage-tolerance of coextruded, bioinspired (“nacre-like”), alumina/nickel compliant-phase ceramics. Scr. Mater. 158, 110–115. (doi:10.1016/j.scriptamat.2018.08.046)

69. Cao SC, Liu J, Zhu L, Li L, Dao M, Lu J, Ritchie RO. 2018 Nature-Inspired Hierarchical Steels. Sci. Rep. 8, 1–7. (doi:10.1038/s41598-018-23358-7)

70. Green KA, Pollock. TM, Harada TE, Howson RC, Reed. JJ, Chirra., Walston. S. 2004 Superalloys 2004. TMS (The Minerals, Metals and Materials Society).

71. Mitchell RJ, Lemsky JA, Ramanathan R, Li HY, Perkins KM, Connor LD. 2008 Process Development & Microstructure & Mechanical Property Evaluation of a Dual Microstructure Heat Treated Advanced Nickel Disc Alloy. In Superalloys 2008 (eds R Reed, K Green, P Caron, T Gabb, M Fahrmann, E Huron, SA Woodard), pp. 347–356.

72. Mitchell RJ. 2010 Polycrystalline Nickel-Based Superalloys: Processing, Performance, and Application. Encycl. Aerosp. Eng., 1–12. (doi:10.1002/9780470686652.eae217)

73. Loh GH, Pei E, Harrison D, Monzón MD. 2018 An overview of functionally graded additive manufacturing. Addit. Manuf. 23, 34–44. (doi:10.1016/j.addma.2018.06.023)

